# Optical control of cardiac rhythm by *in vivo* photoactivation of an ERG channel peptide inhibitor

**DOI:** 10.1101/2023.03.23.534038

**Authors:** Jérôme Montnach, Hugo Millet, Antoine Persello, Hervé Meudal, Stephan De Waard, Pietro Mesrica, Barbara Ribeiro, Jérémie Richard, Agnès Hivonnait, Agnès Tessier, Benjamin Lauzier, Flavien Charpentier, Matteo E. Mangoni, Céline Landon, Chris Jopling, Michel De Waard

## Abstract

**RATIONALE:** Cardiac rhythm, conduction and synchronization of electrical activity require the coordinated action of different types of ion channels that differ according to transmural and regional specificities. Classical pharmacology affects these ion channels in a non-regionalized way which explains why treating arrhythmias, that often occur in specific foci, has often limited efficacy in addition to negative side-effects on non-targeted organs. Photopharmacology is an emergent technology that has the potential to counteract all the negative aspects of classical pharmacology by restricting drug activity in a spatio-temporal manner.

**OBJECTIVE:** We tested the potential of photopharmacology in specifically regulating heart activity by using a caged derivative of a natural peptide inhibitor of the ERG channel, BeKm1. The peptide was uncaged and activity monitored *in vitro* on a cell line expressing the hERG channel, on human cardiomyocytes derived from iPS cells, and *ex vivo* and *in vivo* on zebrafish larvae and rat hearts.

**METHODS AND RESULTS:** Caged BeKm-1 is inactive and fully active upon uncaging. Uncaging of the peptide on human iPS-derived cardiomyocytes enlarges the action potential duration and triggers arrhythmias. Uncaging also triggers bradycardia and disturbs cardiac conduction within the atria in perfused rat hearts upon illumination. The potency of photopharmacology for cardiac electrical modulation was further validated in zebrafish larvae where illumination of the caged compound induces bradycardia and atrio-ventricular desynchrony. Finally, in anesthetized rats, illumination of the caged peptide in the right atria, containing the sino-atrial node, leads to bradycardia without arrhythmia.

**CONCLUSIONS:** This report demonstrates that photopharmacology, using the caged peptide strategy, can be used for dynamically regulating cardiac electrical activity *in vivo* and that spatial illumination restriction can dissociate the bradycardic effect from the arrhythmic one. The technology is applicable to all kinds of cardiac ion channels and regions of interest to create arrhythmogenic models or investigate new clinical applications.

---

Sudden cardiac death is the most frequent cause of cardiovascular mortality in the world (15 to 20% of all deaths and almost 50% of cardiovascular mortality)^1^ and represents a major social burden and public health concern. In addition to atrial fibrillation, which is the most frequent cardiac arrhythmia^2^, ventricular arrhythmias represent more than 75% of sudden death cases^1^. Whether the origin is genetic, drug-induced or secondary to structural defects (hypertrophy, myocardial infarction, …), cardiac arrhythmias always involve ion channel dysfunction. Clinical recommendations consist of the use of implantable pacemakers or cardioverter–defibrillators (ICD)^3^ and/or β-blockers and anti-arrhythmic administration or catheter ablation to treat arrhythmias^4^. Clinical usage of anti-arrhythmic agents is still limited due to their narrow therapeutic window between the ineffective and the paradoxical pro-arrhythmic dose^5^. In addition, conventional molecules have serious extracardiac adverse effects that may lead to increased mortality^6^. All these approaches for the management of arrhythmias, while being extremely beneficial to society, are still not selective enough to avoid a number of worrisome secondary effects. Therefore, there is clinical interest for the development and validation of new innovative therapeutic approaches.

For years, pharmaceutical companies have attempted to discover new compounds active on cardiac ion channels for the treatment of arrhythmias that would lack several of the secondary effects of the existing drugs^7^. Because of the overlap in tissue expression of a given ion channel target, this classical pharmacology approach remains unsuccessful and has not brought any competitive advantage to the existing clinical drugs. Another reason for the failure to develop adapted cardiac pharmacology is that arrhythmias need to be treated at the time of occurrence and not before or after which is hardly compatible with the actual modes of administration. Hence, cardiac pharmacology has to be reinvented by additional pioneering features. By allowing a spatio-temporal control of ion channel activity by light, both optogenetics and photopharmacology appear as promising approaches to equally solve the selectivity and temporal issues in treating arrhythmias. Impressive and convincing reports have illustrated the blooming power of optogenetics for the treatment of arrhythmias^8–12^. Nevertheless, while this approach is powerful, it suffers from one major drawback which is the need to genetically modify the heart in order to express photosensitive ion channels, which is irreconcilable so far with clinical applications. In contrast, photopharmacology does not necessarily require any genetic manipulation which opens interesting avenues for its application to the clinics. Surprisingly, this technology has been poorly investigated for regulating cardiac activity in spite of its successful application for vision restoration^13^ or nociception^14^. In a few cases, cardiac photopharmacology has shown its promises by using photoswitchable chemical drugs. However, the principle of photoswitching of a drug – toggling between an active and an inactive conformation depending on light wavelength -, while being intellectually attractive, has shown moderate efficacy^15–17^. The reason for this failure to deliver the promise is that the photoswitches have incomplete efficacies (a large proportion of the injected compound is already active at a wavelength that is supposed to render it inactive) and the compounds themselves have low affinity requiring high concentrations to be injected *in vivo*.

Recently, we have developed a new approach in photopharmacology that takes advantage of the exquisite *in vivo* properties of natural peptides targeting ion channels^18^. This technology is based on the use of caged peptides allowing the release of the active principle by light in a spatio-temporal manner. It leads to the production of the active compound exclusively in the tissue area of interest and at a concentration that requires minute amounts of the compound. We have first validated this principle by targeting the neuromuscular junction where paralysis of a single muscle could be achieved by the photoactivation of a compound inhibiting the Na_v_1.6 channel^18^. In this report, we now investigate for the first time how this technology can be harnessed for the photocontrol of heart activity by ion channel modulation. For this first proof-of-principle demonstration, we first sought to test this technology for its ability to regulate heart rhythm. To achieve this, we chose to target the ERG channel (K_v_11.1) with a natural peptide, BeKm-1, a 36 amino-acid peptide with three disulfide bridges originally isolated from the venom of *Buthus eupeus* scorpion^19^, which has both an exquisite selectivity and nanomolar potency for K_v_11.1^19–23^. In healthy tissues, this channel is expressed in the sino-atrial node (SAN) where it controls, along with other channel types, pacemaker action potential and rhythmicity ^24–30^. Depending on species, it can also be expressed in the atria and ventricles (human and zebrafish, but not rodents) ^31, 32^. Here we describe the chemical production and functional validation of a caged BeKm-1 peptide and illustrate how this new compound permits optical control of cardiac ERG channels in several models upon light illumination *in vitro*, *ex vivo* and *in vivo*. By allowing an optical control of ERG channel activity in integrated models, our study brings the first evidence that ion channel activity control by peptide-based photopharmacology has the potential to fine-tune cardiac activity *in vivo.* This technology should ultimately help understand the functional role of ion channels to cardiac pacemaker activity or arrhythmogenesis.

## Methods

### Ethics oversight

All animal care and experimental procedures were performed in the animal facilities that have been accredited by the French Ministry of Agriculture. All animals were exposed to 12-h light/dark (light, 8:00 h to 20:00 h) in a thermostatically controlled room with free access to food and water. The experimental procedures were approved by the regional ethic committees (CEEA-006 Pays de la Loire and CEEA-036 Languedoc Roussillon, France) and authorized by the French Ministry of National Education (APAFIS #34541-2022010310194375 and 2017010310594939), Higher Education and Research according to the Directive 2010/63/EU of the European Union. Zebrafish larvae (*Danio rerio*, wild-type AB strain) were conducted in accordance with local approval (APAFIS#4054-2016021116464098 v5) and the European Communities council directive 2010/63/EU

### Data availability

The authors declare that all supporting data are available within the article and in its Supplemental Material. The data that support the findings of this study and analytical tools are available upon request.

### Molecular modelling

To interpret how modifications in BeKm-1 peptide sequences are linked to alterations in channel interaction, molecular models of the interactions were built. The previously published docking model^33^ of BeKm-1 (pdb 1J5J) onto the hERG channel (pdb 5VA1) was used as a template for i) illustrating BeKm-1 interaction on hERG, ii) deciding which residue of BeKm-1 (K^18^) is a good candidate for caging, and iii) highlighting the Van-der Waals (VdW) clashes induced by the photosensitive Nvoc protecting group (cage) on the lateral chain of K^18^. VdW repulsion forces between caged-BeKm-1 and hERG were calculated using the “Show bumps” plugin, implemented in PyMOL (http://www.pymolwiki.org/index.php/Show_bumps).

### Chemical syntheses of BeKm-1 and caged-BeKm-1

All peptides were assembled stepwise using Fmoc-based Solid Phase Peptide Synthesis (SPPS) on a PTI Symphony synthesizer at a 0.05 mmol or 0.1 mmol scale on 2-chlorotrityl chloride polystyrene resin (substitution approximately 1.6 mmol/g). For caged BeKm-1, Fmoc-L-Lysine residues at position 18 were replaced by Fmoc-L-Lys(Nvoc)-OH (Iris Biotech, Marktredwitz, Germany) during assembly. The Fmoc protecting group was removed using 20% piperidine in DMF and free amine was coupled using tenfold excess of Fmoc amino acids and HCTU/DIEA activation in NMP/DMF (3x15 min). Linear peptides were de-protected and cleaved from the resin with TFA/H_2_O/1,3-dimethoxybenzene(DMB)/TIS/2,2′- (Ethylenedioxy)diethanethiol(DODT) 85.1/5/2.5/3.7/3.7 (vol.), then precipitated out in cold diethyl ether. The resulting white solids were washed twice with diethyl ether, re-suspended in H_2_O/acetonitrile and freeze dried to afford crude linear peptide. Oxidative folding of the crude linear caged-BeKm-1 was successfully conducted at room temperature (RT) in the conditions optimized for the BeKm-1 using a peptide concentration of 0.1 mg/mL in a 0.1 M Tris buffer at pH 8.0 containing 10% DMSO. Purification of the peptides were performed on Chromolith®HighResolution RP-18e column (100 x 4.6 mm) using a 5-65% buffer B run in 14 min at 2 mL/min. Solvent system: A: H_2_O 0.1%TFA and B: MeCN 0.1%TFA.

### LC-ESI-QTOF MS analyses

LC-ESI-MS data were acquired with an Accurate mass QTOF 6530 (Agilent) coupled to an Agilent 1290 UPLC system. If required, separation of the peptide sample (5 µL, approx. 10 µg/mL) was done on an Ascentis express 90 Å C18 column (2.0 μm, 2.1 mm ID × 100 mm L, Supelco) heated at 70°C at a flow rate of 0.5 mL/min and with a 2-70% buffer B/A gradient over 11 min (buffer A: H_2_O/formic acid, 99.9/0.1 (v/v); buffer B: ACN/formic acid, 99.9/0.1 (v/v)). If not required, the sample was directly injected at 0.6 mL/min flow rate in H_2_O/MeCN +0.1% formic acid. Acquisition was done in the positive mode with a Dual ESI source, within a *m/z* 100-1700 mass range and analyzed *via* the Agilent MassHunter software version 10.0. Source parameters were set as follows: capillary voltage, 4 kV; fragmentor, 100 V; gas temperature, 350°C; drying gas flow, 12 L/min; nebulizer 50 psig. The observed monoisotopic masses of the peptides were as follow: BeKm-1: 4088.79 Da (theoretical 4088.78); caged BeKm-1: 4327.85 Da (theoretical: 4327.81).

### Analytical RP-HPLC analyses

Analytical RP-HPLC was performed to monitor Nvoc deprotection of caged BeKm-1. Using an SPD M20-A system (Shimadzu) with a Luna OmegaPS C18 column (4.6 x 250 mm, 5 μm, 100 Å), 20 µL of peptide (corresponding to 7 µg of material) was loaded and a 5-60% acetonitrile gradient (0.1% TFA v/v) was applied over 35 min at RT to detect analytes at 214 nm. Illumination of samples was performed at 365 nm for different times (between 1 sec and 30 min) at 41.8 mW/cm^2^ or for different power of illumination (as specified in the Result section) for 10 min using a CoolLED pE4000 light source (CoolLED, UK). Flash energy has been measured using a highly sensitive thermal power head (S401C, ThorLabs). Coelution of uncaged BeKm-1 and non-caged BeKm-1 was performed using a 50:50 ratio for both compounds.

### NMR spectrometry

Caged BeKm-1 was illuminated for 30 min at 100% power (42 mW/cm^2^) using a CoolLED pE4000 light source (CoolLED, UK) to produce fully uncaged BeKm-1. For NMR spectra acquisitions, three 200-μL solutions were prepared in 3-mm NMR tubes containing i) caged BeKm-1 (500 µM), ii) uncaged BeKm-1 (500 µM) or iii) non-caged BeKm-1 (500 µM), respectively. For each sample, a set of 2D homonuclear (80 ms TOCSY and 160 ms NOESY) and heteronuclear (^13^C-HSQC and ^15^N-HSQC, natural abundancy) spectra were acquired on a BRUKER 700 MHz NMR spectrometer equipped with a 5-mm TCI cryoprobe, at 298K. Processing and analyses were performed with Bruker’s TopSpin3.6 and CcpNMR2.1.5 programs. Spectra were drawn with CcpNMR program, and 3D structure were drawn with the PyMOL2.3 software (The PyMOL Molecular Graphics System, Schrödinger, LLC).

### Cell cultures

HEK293 cells stably expressing hERG (NM_000238) channels were cultured in Dulbecco’s Modified Eagle’s Medium (DMEM) supplemented with 10% fetal bovine serum, 1 mM pyruvic acid, 4.5 g/L glucose, 4 mM glutamine, 800 µg/mL G418, 10 U/mL penicillin and 10 μg/mL streptomycin (Gibco, Grand Island, NY). hERG cells were incubated at 37°C in a 5% CO_2_ atmosphere. For electrophysiological recordings, cells were detached with trypsin and floating single cells were diluted (∼300,000 cells/mL) in medium containing (in mM): 140 NaCl, 4 KCl, 2 CaCl_2_, 1 MgCl_2_, 5 glucose and 10 HEPES (pH 7.4, osmolarity 298 mOsm).

### Automated patch-clamp recordings and pharmacological studies

Whole-cell recordings were used to investigate the effects of BeKm-1 analogues on HEK293 cells expressing hERG channels. Automated patch-clamp recordings were performed using the SyncroPatch 384PE from Nanion (München, Germany). Chips with single-hole medium resistance of 4.52 ± 0.08 MΩ (n=384) were used for recordings. Pulse generation and data collection were performed with the PatchControl384 v1.5.2 software (Nanion) and the Biomek v1.0 interface (Beckman Coulter). Whole-cell recordings were conducted according to the recommended procedures of Nanion. Cells were stored in a cell hotel reservoir at 10°C with shaking speed at 200 RPM. After initiating the experiment, cell catching, sealing, whole-cell formation, liquid application, recording, and data acquisition were all performed sequentially and automatically. Intracellular solution contained (in mM): 10 KCl, 110 KF, 10 NaCl, 10 EGTA and 10 HEPES (pH 7.2, osmolarity 280 mOsm). Extracellular solution contained (in mM): 140 NaCl, 4 KCl, 2.48 CaCl_2_, 1.69 MgCl_2_, 5 glucose and 10 HEPES (pH 7.4, osmolarity 298 mOsm). Cells were maintained at a holding potential of −80 mV and BeKm-1 analogues were tested during a 50-ms test potential at +60 mV following a first activation step of 1000-ms at +60 mV and a 10-ms step at -120 mV to recover from inactivation with a pulse every 8 sec. The pharmacology of caged BeKm-1, as well as the efficiency of the released product to block hERG channels was studied using combined automated patch-clamp and UV illumination. After 2 min of control period, caged BeKm-1 was added to the external buffer and effects were recorded for 100 sec prior to a 250-sec duration illumination for photocleavage induction of the caged compound. Different wavelengths, illumination powers and durations were used as specified in figure legends.

### *Ex vivo* illumination on Langendorff-perfused hearts

Ten-week-old rats were heparinized (heparin sodium, 0.5 U/g intraperitoneal (i.p.)) and euthanized by i.p. injection of lethal dose of pentobarbital. Hearts were quickly removed and rinsed in a modified cold (4°C) Krebs solution containing (in mM): 116 NaCl, 5 KCl, 1.1 MgSO_4_, 0.35 NaH_2_PO_4_, 27 NaHCO_3_, 10 glucose, pH 7.4. Hearts were rapidly cannulated and perfused in a retrograde fashion through an aortic cannula with warm (37-38°C) oxygenated modified Krebs solution supplemented with (in mM): 1.8 CaCl_2_, 0.2 pyruvate, 1 lactate at a constant flow rate of 12 mL/min without recirculation. Once connected to the Langendorff perfusion system, hearts were perfused with control solution for 10 min to stabilize their activities. Hearts were electrophysiologically assessed using a 36-electrode arrays flexible MEA system (EcoFlexMEA 36; MultiChannel Systems, Reutlingen, Germany). Data were sampled at 10 kHz per channel with simultaneous data acquisition using Cardio 2D. Caged BeKm-1 (250 nM) was perfused and illumination at 365 nm at 75 mW/cm^2^ applied at 3 min using a CoolLED pE4000 light source (CoolLED, UK) for 10 min. Data were analyzed using Cardio 2D+ (MultiChannel Systems, Reutlingen, Germany).

### Patch-clamp recordings of sino-atrial node cells

Wild-type (Charles River laboratory) mice were bred and maintained under the C57Bl/6J genetic background. Sino-atrial node (SAN) cells were isolated from C57Bl/6J mice as previously described^28, 34^. SAN cells were harvested in custom-made chambers with glass bottoms for cell attachment and superfused with Tyrode solution warmed at 36°C before recording. Action potentials (APs) were measured by the perforated patch-clamp technique and recorded using a Multiclamp 700A patch-clamp amplifier connected to a Digidata 1550B interface (Molecular Devices). Electrodes had a resistance of 3-4 MΩ when filled with a solution containing (in mM): 80 K-aspartate, 50 KCl, 1 MgCl_2_, 2 CaCl_2_, 5 EGTA, 5 HEPES, and 3 ATP-Na (adjusted to pH 7.2 with KOH). Perforated patch-clamp was performed by adding 30 µM β-escin to the intracellular solution. Perfusion of pre-warmed 50 nM BeKm-1 solution was achieved by using a multi-MPRE8 heating pen (Cell Micro Controls). AP firing rate and maximal diastolic potential (MDP) were measured as previously described^35^.

### Action potential recordings in hiPS-CMs

Single-cell dissociated hiPS-CMs were plated at low density on matrigel-coated 35- mm Petri dishes (Nunc) and electrophysiological experiments were performed 12-14 days after plating. APs were recorded at 37°C in the current-clamp mode using the dynamic patch-clamp method to electronically mimic expression of the inward rectifier potassium current, I_K1_^23, 36, 37^. Two pA/pF *in silico* I_K1_ density (maximum density on the current-voltage curve at -72 mV) was sufficient to bring the resting membrane potential (RMP) to -80 mV ± 5 mV. Cells were depolarized every 1000 ms with a 30-35 pA/pF current density 1-ms pulse. Both cell pacing and I_K1_ injection were driven by a custom-made software and a National Instrument A/D converter (NI PCI-6221) connected to the current command of the amplifier. Cells were superfused with extracellular solution containing (in mM): 140 NaCl, 4 KCl, 0.5 MgCl_2_, 1 CaCl_2_, 10 HEPES, 10 glucose, pH 7.4 (NaOH). Low resistance borosilicate glass pipettes (2-3 MΩ; Sutter Instruments) were filled with a solution containing (in mM): 125 K-gluconate, 5 NaCl, 20 KCl, 5 HEPES, pH 7.2 (KOH), supplemented with 0.22 mg/mL amphotericin-B. Stimulation and data recordings were performed using an Axopatch 200B amplifier controlled by Axon pClamp 10.6 software through an A/D converter (Digidata 1440A, Molecular Devices). The data were analyzed using home-made R scripts. Various AP characteristics were examined, including maximum diastolic potential (MDP), AP overshoot (*i.e.* amplitude of the AP over 0 mV), maximal upstroke velocity (dV/dt_max_), and AP duration at different times of repolarization (APD). For each analysis, parameters of APs over 30 sec were averaged to reduce intracellular AP variability.

### Electrical field potential measurements of hiPS-CMs

The electrical field potentials (EFPs) of hiPS-CMs were evaluated with a CardioExcyte96 system. After dissociation, hiPS-CMs were seeded on NSP-96 Sensor Plates (Nanion Technologies, Munich, Germany) at a density of 40,000 viable cells per well. Sensor plates were previously coated with 10 µg/mL fibronectin for 1 h at 37°C. The cells were maintained at 37°C, and the medium was replaced every 2 days for 1 week. On the day of the experiment, the whole medium was replaced 4 h prior to experiments, and cells were monitored with the CardioExcyte96 system to ensure the full stabilization. Once this was achieved, caged BeKm-1, followed by illumination, could be applied. Ten EFPs were analyzed as control prior to addition of caged BeKm-1 (or control medium), while several other EFPs were recorded before and after illumination of the wells at 365 nm.

### Electrocardiography

Wild-type C57Bl/6J mice were anesthetized with 2% isoflurane *via* a nose cone (0.3 L/min following induction in a chamber containing isoflurane 4-5%). Rectal temperature was monitored continuously and maintained at 37-38°C using a heat pad. Six leads ECGs were recorded with 25-gauge subcutaneous electrodes on a computer through an analogic-digital converter (IOX 1.585, EMKA Technologies, France). ECG channels were filtered between 0.5 and 250 Hz. ECGs were recorded during 10 min after atropine sulfate/propranolol injection (0.5 mg/kg and 1 mg/kg, respectively i.p.; Control group) or after atropine sulfate/propranolol and BeKm-1 (0.5 mg/kg, i.p.; BeKm-1 group) or after atropine sulfate/propranolol and caged BeKm-1 (5 mg/kg, i.p.; Caged BeKm-1 group). Atropine sulfate and propranolol were used to block the autonomic nervous system. Reported measurements of PQRST complexes were averaged by 10-s sections in lead I (ECG Auto v3.5.5.28, EMKA Technologies). Standard criteria were used for interval measurements. The end of the T wave was defined as the point at which the slow component returned to the isoelectric line. Area of T wave is measured between S wave and the end of T wave. QT intervals were corrected for heart rate using the formula, QTc=QT/(RR/100)^1/2^ with QT and RR measured in ms ^38^.

### Zebrafish larvae heart rate tracking

Zebrafish larvae (*Danio rerio*, wild-type AB strain) were maintained under standardized conditions. One hundred twenty hours post fertilization (hpf) larvae were anaesthetized using tricaine methane sulfonate (MS222, Sigma-Aldrich). Larvae were injected (microinjector-World Precision Instruments) in the pericardium with 1 nL of either vehicle (Texas Red) or a 50-μM solution of caged BeKm-1. After 10 min of tracking, larvae were exposed to 365 nm light (Prismatix) for 5 min. Videos were acquired at 200 fps and analyzed using Fiji software in order to provide the cardiac rhythm of both atrium and ventricle. The brightness signal of the analyzed region was computed as pixel variations between frames and a Fourier transform is calculated using the Fast Fourier Transform (FFT) algorithm to translate the signal from the time domain to the frequency domain. The frequency of the first FFT peak was transformed to beats per minute.

### *In vivo* photoactivation of caged BeKm-1 in rat hearts

Ten-week-old Wistar Han rats were used to evaluate the impact of caged BeKm-1 photoactivation on the cardiac rhythm of anesthetized rats. The procedure was performed under anesthesia using isoflurane. Rats were intubated with a 16-gauge catheter and ventilated using Harvard Apparatus model 683 (Harvard Apparatus, Holliston, MA). The right atria were exposed *via* right thoracotomy in the 2^nd^ intercostal space. The thymus was excised to prevent shadow on the atria during illumination. Six-lead ECGs were recorded with 25-gauge subcutaneous electrodes on a computer through an analogic-digital converter (IOX 1.585, EMKA Technologies, France). ECG channels were filtered between 0.5 and 250 Hz. ECGs were recorded right after thoracotomy (Control group). Illumination of the right atrium (365 nm, 75 mW/cm²) was performed using the CoolLED system 3 min after injection of caged BeKm-1 (100 µg in 100 µL) into the dorsal penile vein. Reported RR interval was averaged by 10-sec sections in lead I (ECG Auto v3.5.5.28, EMKA Technologies).

### Statistics and data analyses

Values are represented as mean ± SEM. Significance of illumination tests on automated patch-clamp was tested by performing paired 1-way ANOVA and Bonferroni’s multiple and significance of *in vivo* experiments was tested by performing Friedman test with Dunn’s multiple comparisons test. A p-value lower than 0.05 was considered significant.

## Results

### BeKm-1 induces bradycardia by targeting SAN cells

We have investigated the effects of BeKm-1on heart rate and SAN pacemaker activity in wild-type mice using 6-leads ECGs after injection of 0.5 mg/kg atropine and 1 mg/kg propranolol to concomitantly block parasympathetic and sympathetic modulation of heart rate, respectively (Figure S1a,b). Injection of 0.5 mg/kg of BeKm-1 induced bradycardia evidence by significant increase in RR intervals (133.6 ± 4.8 ms with BeKm-1 *versus* 118.0 ± 3.3 ms in control condition before BeKm-1 injection, n= 5, p<0.05). As previously described for ERG blockers^39^ in mice, corrected QT interval (QTc) was statistically unchanged despite prolongation of QT interval resulting from bradycardia. Other ECG parameters were not altered by injection of BeKm-1 (Table S1). The absence of induction of arrythmia by BeKm-1 was coherent with a minor contribution of ERG channels to cardiac repolarization in mice^40^. We then studied whether the bradycardic effects of BeKm-1 were due to an effect on SAN pacemaker cells. Fifty nM of BeKm-1 induced a significant 2.9-fold reduction of AP rate in isolated SAN cells from adult mice (78.8 ± 16.6 bpm with BeKm-1 *versus* 231.0 ± 11.7 bpm in control condition prior to BeKm-1 application, n= 18, p< 0.0001) (Figure S1c,d). It also shifted the maximal diastolic potential to more positive voltages (-52.6 ± 2.9 mV *versus*-62.4 ± 0.9 mV, n= 18, p< 0.01) and prolonged the action potential duration (APD) both at 70% and 90% of full repolarization (81.7 ± 12.9 ms *versus* 57.2 ± 4.9 ms, n= 11-18, p< 0.05 and 124.4 ± 20.2 ms *versus* 80.8 ± 6.9 ms, n= 11-18, p< 0.05, respectively) (Figure S1e,f). Taken together, these results showed that BeKm-1 induced bradycardia in mice by inhibiting automaticity of SAN pacemaker cells.

### Design and chemical synthesis of caged BeKm-1

To produce an inactive caged BeKm-1 peptide that can be experimentally activated by light, we chose to chemically synthetize BeKm-1 with a photosensitive orthogonal o- nitroveratryloxycarbonyl (Nvoc) (Figure 1a). For the position of the Nvoc group, K^18^ was chosen because it is a key amino-acid for hERG/BeKm-1 interaction based on the published hERG/BeKm-1 complex^33^ (Figure 1b; interaction with G^626^ and F^627^ of the hERG channel, two residues located on two distinct monomers) and according to results from a previous report showing that the K^18^A mutation drastically reduces the affinity of the peptide for hERG^20^. By using this docked hERG/BeKm-1 complex structure^33^, we calculated that addition of the Nvoc group on the lateral chain of K^18^ would induce severe van der Walls steric clashes with S^624^, V^625^, G^626^ and F^627^ of distinct monomers of hERG. The direct effects of Nvoc addition, modifying the K^18^ properties, and the local reorganization of hERG expected to occur to prevent steric clashes, should be sufficient to drastically reduce the affinity of the peptide for hERG (Figure 1c). Next, caged BeKm-1 was assembled stepwise using Fmoc-based solid phase peptide synthesis (Figure 1d). Proper folding of caged peptide was confirmed by the disappearance of the unfolded peptide and formation of one main oxidized compound, slightly shifted on the elution profile to greater water polarity, that was purified. The proper mass of caged BeKm-1 was confirmed with a measured *m/z* ratio of 1082.96 [M+4H]^4+^ corresponding to a peptide of 4327.88 Da (theoretical monoisotopic mass of 4327.81 Da, Figure 1e) suggesting that the Nvoc group was preserved during the entire synthesis process (non-caged BeKm-1 monoisotopic mass of 4091.7 Da for comparison). Using NMR spectrometry, we showed that the presence of the Nvoc group on K^18^ disturbs the chemical environment of residues in proximity with K^18^ (significant shifts for S^10^, Y^11^, V^16^, C^17^, S^19^, R^20^, K^23^, T^24^, N^25^ and F^36^, disappearance of the peaks of Q^12^, F^21^, and G^26^ in the caged form) (Figure 1f,g and Table S2). The NMR data also confirm that the global folding of caged BeKm-1 is not disrupted. Overall, the presence of the Nvoc group on the lateral chain of K^18^, combined to the alterations in space of the lateral chain of residues surrounding the central caged K^18^, should greatly affect the inhibitory activity of caged BeKm-1 on hERG channel. Using HEK293 cells stably-expressing the hERG channel, we illustrate that caged BeKm-1 is unable to inhibit hERG-mediated K^+^ currents at concentrations (between 0.1 and 100 nM) at which non-caged BeKm-1 is highly potent (Figure 1h). For instance, 100 nM of BeKm-1 blocks 93 ± 2% of hERG currents, whereas caged BeKm-1 has no effect at all (0.6 ± 2% of inhibition) on the amplitude of the K^+^ currents (Figure 1h). Concentration-response curves illustrate that caging reduces the potency of the peptide by 2,400-fold (non-caged BeKm-1: IC_50_ = 1.6 nM, n = 46 cells; caged BeKm- 1: IC_50_ = 3,847 nM, n = 94 cells). From these data, it is concluded that caged BeKm-1 is an inert compound at concentrations up to 300 nM.

**Figure 1.**
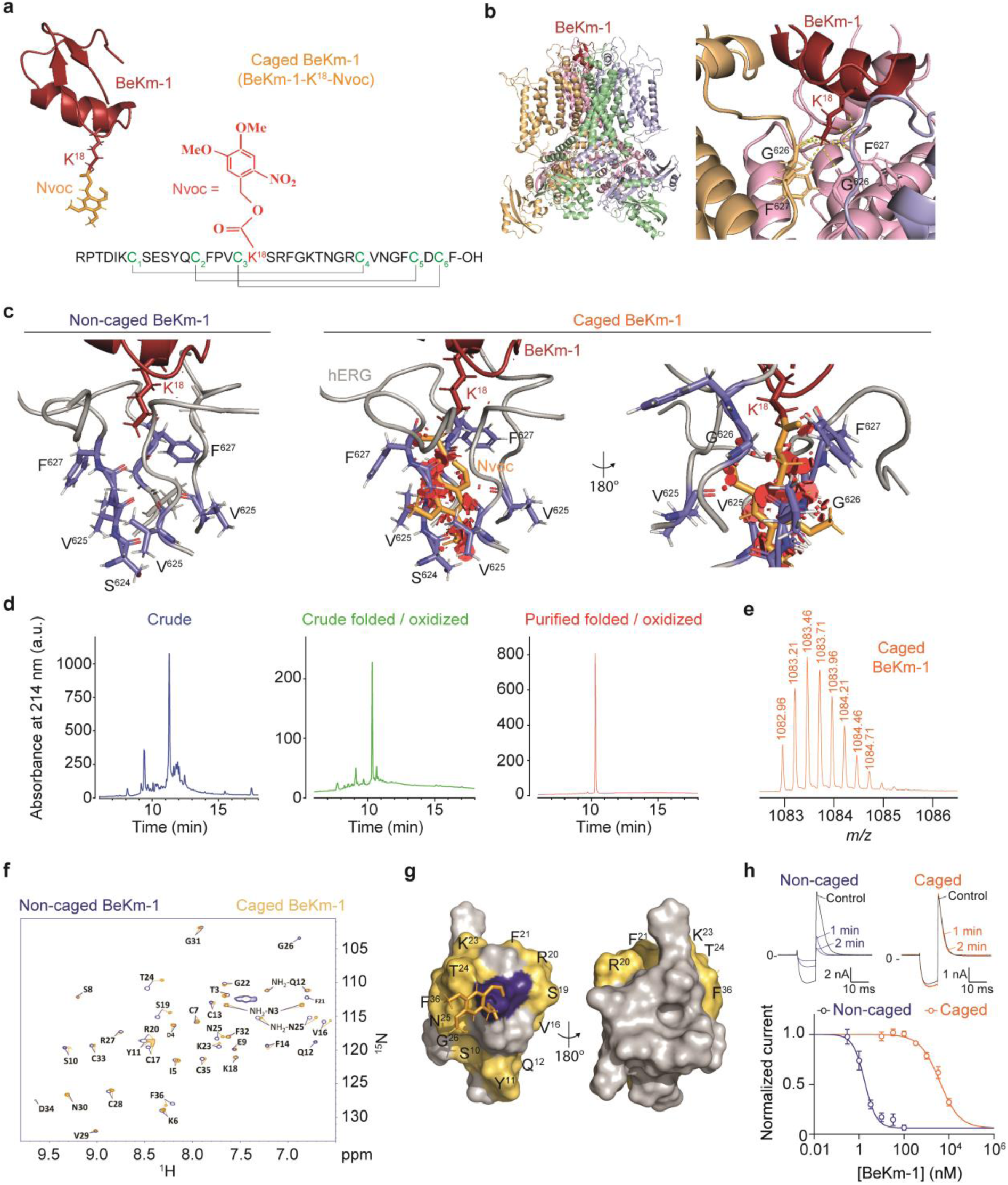
Engineering of photoactivatable caged BeKm-1 to modulate ERG channels. (a) Structure of caged BeKm-1 with K^18^ and Nvoc group reported in red and orange sticks respectively. (b) Docking model of BeKm-1 on hERG channel showing interactions between K^18^ of BeKm and hERG channel. BeKm-1 and hERG structures have been obtained from PDB 1J5J and 5VA1, respectively. (c) Illustration of steric van der Walls clashes (red cylinders) resulting from the addition of the Nvoc protecting group onto K^18^ upon binding of BeKm-1 onto hERG channel in the position as published ^33^ (PDB code 5VA1). (d) RP-HPLC elution profiles of crude, crude folded/oxidized, and purified folded/oxidized caged BeKm-1. (e) LC-ESI QTOF analysis of caged BeKm-1 denotes an exact mass of 1082.96 [M+4H]^4+^. (f) Superimposition of ^15^N-so-Fast-HMQC spectra of the non-caged BeKm-1 (in blue), and of caged BeKm-1 (in orange). (g) Residues showing major chemical shifts modifications in the presence of Nvoc are reported in orange on the surface of the molecule (180°views). K^18^ is highlighted in blue. The Nvoc group is reported in orange sticks. (h) Representative recording of hERG currents elicited at +60 mV following a first activation step of 1000 ms at +60 mV and a 10-ms step at - 120 mV to recover from inactivation illustrating the extent of current block by 100 nM of non-caged BeKm-1 (blue) and caged BeKm-1 (orange). ii) Average dose-response curves for hERG current inhibition by non-caged BeKm-1 (blue) and caged BeKm-1 (orange). The data were fitted according to a Hill equation (non-caged BeKm-1: IC50 = 1.6 nM, n = 46 cells; caged BeKm-1: IC50 = 3847 nM, n = 94 cells). Data are presented as mean ± SEM.

### Photolysis of caged BeKm-1 restores the exact BeKm-1 peptide

We and others previously described that the photolysis of the Nvoc group grafted on an amino-acid is sensitive to UV wavelengths^18, 41, 42^. Compared to non-caged BeKm- 1, caged BeKm-1 presents a shifted elution profile towards less polar properties on RP-HPLC, consistent with more hydrophobic Nvoc properties. Co-elution experiments of non-caged BeKm-1 with fully uncaged BeKm-1 (after illumination of caged BeKm-1 at 365 nm and at 45 mW/cm² for 30 min) results in a single elution peak suggesting that photolysis releases a BeKm-1 peptide with similar elution properties than non-caged BeKm-1 (Figure 2a). The identity between uncaged and non-caged BeKm-1 peptides was further confirmed by mass spectrometry analyses indicating a common molecular mass (uncaged BeKm-1 *m/z* = 1023.2 [M+4H]^4+^ corresponding to a peptide of 4088.88 Da; non-caged BeKm-1 *m/z* = 1023.2 [M+4H]^4+^ corresponding to a peptide of 4088.88 Da) (Figure 2b). Also, NMR analyses demonstrate that the spectra of non-caged and fully uncaged BeKm-1 are perfectly superimposable (Figure 2c). This finding indicates that all residues in close proximity of K^18^ regain the same spatial distribution and chemical environment as in the non-caged BeKm-1. The proper folding, ensured by the exact pattern of disulfide bridges, is totally preserved (Figure 1a). At the functional level, uncaged BeKm-1 inhibits hERG channels with the same efficacy than non-caged BeKm-1. As for non-caged compound, 100 nM uncaged BeKm-1 produces 92 ± 3% block of hERG K^+^ currents (Figure 1h & Figure 2d). Also, the inhibition by uncaged BeKm-1 occurs with a similar concentration response curve as non-caged BeKm-1 (uncaged BeKm-1: IC_50_ = 1.5 nM, n = 180 cells, non-caged BeKm-1: IC_50_ = 1.6 nM, n = 46 cells; Figure 2d). These results validate that complete photolysis of caged BeKm-1 produces a BeKm-1 that is fully identical in structure and function to the non-caged compound. The uncaging process at 365 nm is time- and UV power-dependent (half-uncaging time of 2.8 min at 45 mW/cm² and half-uncaging power of 9 mW/cm²; Figure S2a-d). Nvoc photolysis is reportedly effective at 350 and 365 nm^18, 43^. Nevertheless, we observe that, at least in the frame of its use for natural peptides, photolysis is effective up to 405 nm which is a better wavelength for tissue penetration and less damaging than 365 nm (Figure S2e-f). No photolysis was observed at wavelengths above 405 nm offering a possibility to combine uncaging with dye-based electrophysiological approaches.

**Figure 2.**
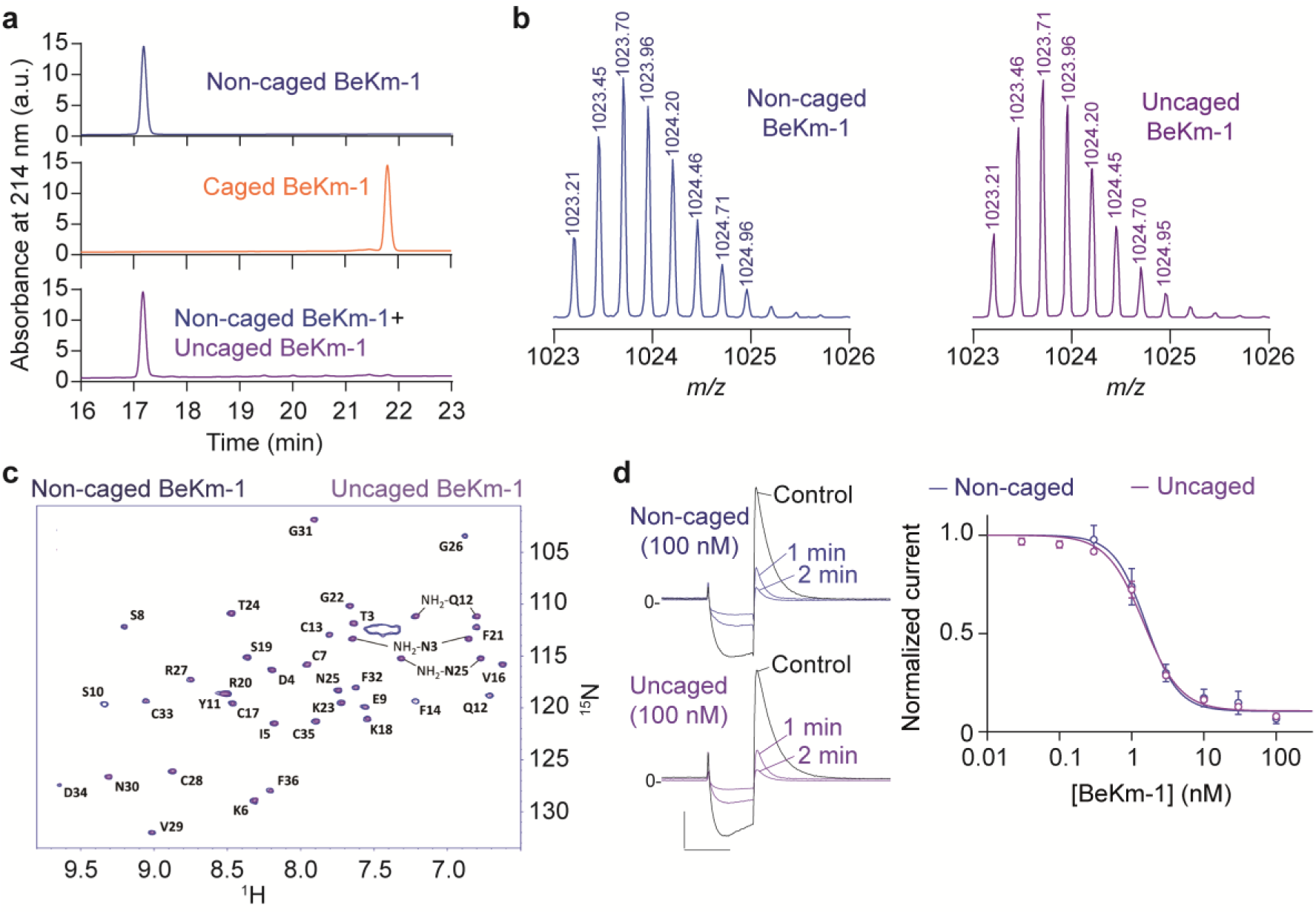
Uncaging of BeKm-1 leads to fully functional BeKm-1 peptide. (a) Analytical RP-HPLC profiles of non-caged BeKm-1 (top), caged BeKm-1 (middle) and 50:50 ratio of non-caged BeKm-1 after uncaging of BeKm-1 (bottom). (b) Mass analyses of non-caged BeKm-1 and uncaged BeKm-1 by LC-ESI QTOF ([M+4H]^4+^ values). (c) Superimposition of ^15^N-so-Fast-HMQC spectra (same area as Fig. 1f) of the non-caged BeKm-1 (in blue), and of the uncaged BeKm-1 after illumination of caged BeKm-1 (in purple). (d) Left: Representative recording of hERG currents elicited at +60 mV following a first activation step of 1000 ms at +60 mV and a 10-ms step at -120 mV to recover from inactivation illustrating the extent of current block by 100 nM of non-caged BeKm-1 (blue) and uncaged BeKm-1 (purple). Scale: 10 ms, 2 nA. Right: Average dose-response curves for hERG current inhibitions by non-caged BeKm-1 (blue) and uncaged BeKm-1 (purple). The data were fitted according to a Hill equation (uncaged BeKm- 1: IC50 = 1.5 nM, n = 180 cells, non-caged BeKm-1: IC50 = 1.6 nM, n = 46 cells). Data are presented as mean ± SEM.

### *In vitro* optical control of hERG activity

Using a high-throughput patch-clamp system, combined with an illumination system, application of 100 nM caged BeKm-1 to HEK293 cells expressing hERG channels has no effect on K^+^ currents, whereas a 3-min illumination of recording wells at 365 nm (45 mW/cm²) produces a progressive and significant inhibition of hERG current in steady-state recording conditions (Figure 3a,b). The kinetics of current block is the combined result of the kinetics of the uncaging process and binding of uncaged peptide onto the channel. Photoactivation of caged 100 nM BeKm-1 leads to a maximal percentage of inhibition of 87.7 ± 1.3% (n=27) in these experimental conditions, whereas the caged peptide has no effect (Figure 3c). We also confirmed that the uncaging process is UV illumination time- and power-dependent at the functional level by demonstrating that the extent of hERG inhibition increases if illumination time or power are increased (Figure S3). We next investigated the effect of photoactivation of caged BeKm-1 on human iPS-derived cardiomyocytes, dynamically clamped at a resting potential of -80 mV and paced at a cycle length of 1,000 ms. As control, we illustrate that illumination of the human iPS-derived cardiomyocytes at 365 nm wavelength has no impact on APD (Figure 3d). Also, application of 200 nM caged BeKm-1 has no effect on APD at 90% of full repolarization (APD_90_: 279.1 ± 47.2 ms *versus* 276.7 ± 44.9 ms, n= 15), whereas photoactivation of the peptide by cell illumination at 365 nm induces a significant average 1.3-fold increase in APD_90_ duration (358.4 ± 55.7 ms *versus* 279.1

**Figure 3.**
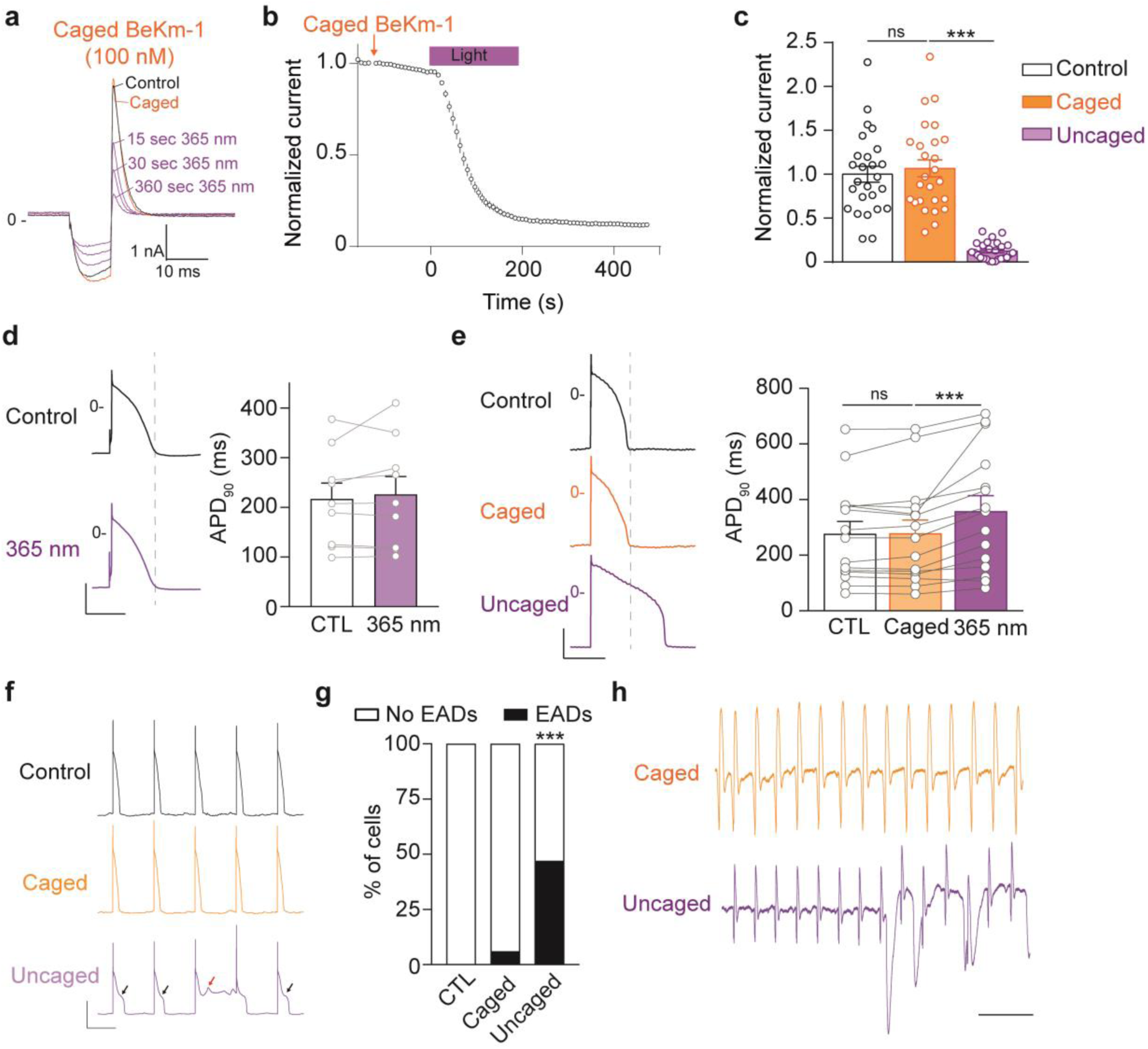
Properties of caged BeKm-1 photolysis. (a) Representative recordings of hERG current in control (black), after incubation with 100 nM of caged BeKm-1 (orange) and after various illumination times (purple). Scale bars: 10 ms, 1 nA. (b) Average normalized time course of hERG current inhibition before, during and following light application (n = 26 cells). (c) Average normalized current at steady-state in control (dark), caged (orange) and uncaged (purple) conditions (n=26, ***p<0.001, two-sided repeated measures 1-way ANOVA followed by Bonferroni’s post-test.). (d) Left: Representative action potentials obtained from IK1- clamped hiPS-derived cardiomyocytes before, during and after 365 nm illumination. Right: Quantification of illumination effects on action potential duration at 90% of full repolarization (n = 9 cells, p=ns). Scale bar: 200 ms and 50 mV. (e) Left: Representative action potentials obtained from IK1-clamped hiPS-derived cardiomyocytes in control condition, after perfusion with 200 nM of caged BeKm-1 and after 365 nm illumination. Right: Quantification of action potential duration at 90% of repolarization (caged BeKm-1: 279.1 ± 47.2 ms *versus* Control: 276.7 ± 44.9 ms; n= 15, p=ns; Uncaged BeKm-1: 358.4 ± 55.7 ms *versus* caged BeKm-1: 279.1 ± 47.2 ms; n= 15, ***p<0.001). Scale bar: 200 ms and 50 mV. (f) Representative AP trains showing the occurrence of EADs after BeKm-1 uncaging in hiPS-derived cardiomyocytes. (g) Percentages of hiPS-derived cardiomyocytes displaying EADs in each condition. (n = 15 cells). Fisher test ***, p<0.001. (h) Representative electrical field potential (EFP) recorded from a layer of hiPS-derived cardiomyocytes incubated with 100 nM caged BeKm-1 (orange) and after illumination at 365 nm (purple). Data are presented as mean ± SEM.

± 47.2 ms, n= 15, p<0.001) (Figure 3e) without affecting other action potential parameters (Table S3). Uncaging of BeKm-1 is also prone to induce early-afterdepolarization in 47% of the cells (Figure 3f-g). In a layer of spontaneously beating hiPS-derived cardiomyocytes, we report that uncaging of BeKm-1 induces slowing of automaticity and dysrhythmia (Figure 3h), which is coherent with earlier observations using non-caged BeKm-1^23^. These results demonstrate the ability to control hERG currents by UV illumination in several cell models.

### Spatio-temporal control of atrial ERG channels in isolated rat heart

To assess if caged BeKm-1 can be used for a spatio-temporal control of hERG channels in cardiac tissue, we evaluated the effect of photoactivation of caged BeKm- 1 in Langendorff-perfused rat heart. By recording the electrical activity of the right atrium using a 36-electrodes array (Figure 4a), we first validated that perfusion of 100 nM BeKm-1 induces a significant 1.5-fold reduction in heart rate compared to the control period (visualized by an increase in inter-beat interval from 217.2 ± 12.7 ms to 316.4 ± 33.9 ms, n= 8, p<0.05, Figure 4b,c). Activation maps indicate that BeKm-1 disrupts atrial conduction and leads to heterogeneity in conduction velocity within the recording area as witnessed by the beat-to-beat variability (Figure 4d). Next, we tested the effect of the perfusion of 250 nM caged BeKm-1 in Langendorff-perfused rat hearts, a concentration that should produce minimal effects on ERG activity (Figure 1h). As expected, such a concentration had indeed minimal effects on RR interval (Figure 4e,f). In contrast, if the right atrium was illuminated 3 min after application of caged BeKm-1 (Figure 4g), then bradycardia was significantly enhanced as witnessed by the increase in RR interval (uncaged BeKm-1: +25.7 ± 2.9 ms *versus* +8.6 ± 2.5 ms for caged BeKm- 1, n= 6, p<0.05) (Figure 4e,f). Activation maps in the illuminated area highlight that caged BeKm-1 does not disrupt atrial conduction, whereas illumination at 365 nm of the right atria disrupts atrial conduction and leads to heterogeneity in conduction velocity within the illuminated area (Figure 4h). In conclusion, these data indicate that illumination of the right atria liberates enough BeKm-1 in the illumination spotlight to impact heart rhythm and to disrupts atrial conduction. Those results indicate that caged BeKm-1 is photoactivatable in an *ex vivo* perfused adult rat heart.

**Figure 4.**
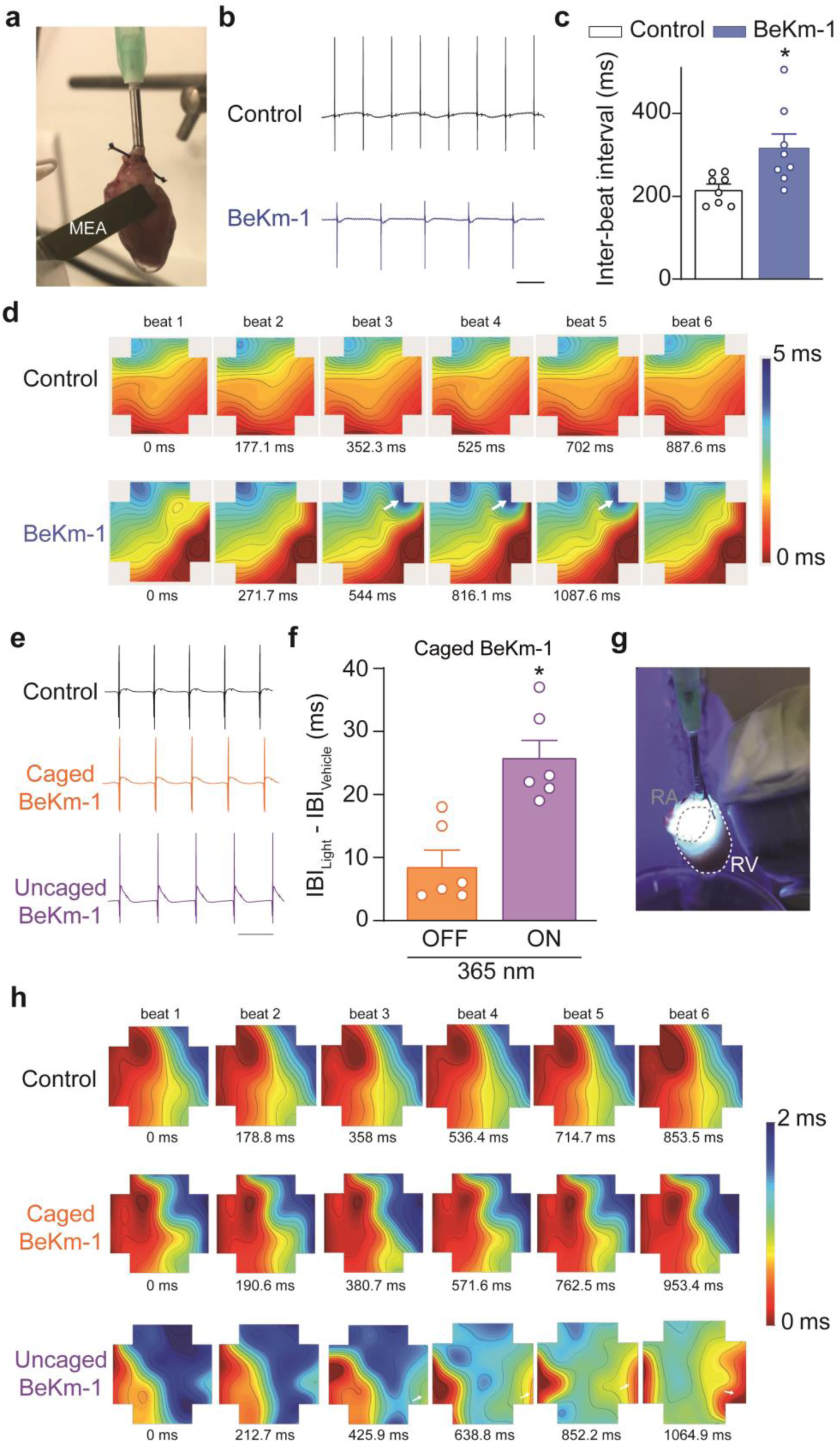
Light-induced inhibition of ERG channels in rat perfused-heart. (a) Multi-electrode array (MEA) setup for recording of electrical right atrium activity, (b) Representative MEA traces obtained in control (black) or after perfusion of 100 nM BeKm-1 during 10 min (blue). Scale = 250 ms. The changes in electrical axis of the recorded MEA area is due to effects of BeKm-1 on cardiac conduction as seen in (d). (c) Quantification of BeKm-1 effects on inter-beat interval (Control: 217.2 ± 12.7 ms *versus* BeKm-1: 316.4 ± 33.9 ms; n= 8 hearts, *, p<0.05). (d) Representative activation maps obtained before and after 10-min perfusion of 100 nM BeKm-1. Colors indicate activation relative to the physiological pacing site according to the scale. Isochrones: 2 ms. Arrows indicate areas with abnormal conduction. (e) Representative MEA traces obtained in control (black), after perfusion of 250 nM caged BeKm- 1 (blue) and after uncaging (purple). Scale = 200 ms. (f) Quantification of inter-beat interval under before and after illumination in caged BeKm-1 perfused hearts (n = 6 hearts, *p<0.05). (g) Photograph showing illumination area focus on the right atrium of the perfused heart. (h) Representative activation maps obtained before and after 10-min perfusion of 250 nM caged BeKm-1 and after uncaging. Colors indicate activation relative to the physiological pacing site according to the scale. Isochrones: 2 ms. Arrows indicate beat-to-beat heterogeneity.

### *In vivo* optical control of ERG activity in zebrafish larvae

After having demonstrated *in vitro* and *ex vivo* the ability to optically control ERG channel activity by caged BeKm-1, we next studied the potential of photoactivation to interfere with cardiac rhythm *in vivo*. We first turned towards the zebrafish larvae, a translucid model frequently used to study the modulation of ERG and convenient in photopharmacology experiments^18^ and optogenetics^44^. zERG being an ortholog ion channel of hERG, we first checked whether BeKm-1 modulates zERG. Injection of 2 nL of 50 µM BeKm-1 in zebrafish larvae produces a significant atrial bradycardia compared to larvae injected with control solution (inter-beat interval of 530.7 ± 27.9 ms in the presence of BeKm-1 *versus* 420.2 ± 13.5 ms in control, n= 10, p<0.01) (Figure 5a, left panel). A similar bradycardia was observed at the ventricular level (Figure 5a, right panel). This bradycardia is associated with a higher variability of the beating rate both at the atrial and ventricular level (Figure 5b). Also, BeKm-1 induces episodes of atrio-ventricular desynchronization in 6/10 larvae (Figure 5c, Movie S1-2). Injection of 2 nL of 50 µM caged BeKm-1 into zebrafish larvae prior to illumination has no effect on inter-beat interval (379.7 ± 13.8 ms with caged BeKm-1 *versus* 420.2 ± 13.6 ms in control, n= 10, Figure 5d). However, illumination (365 nm, 3 min) of the larvae induces a phenotype similar to the one produced by injection of non-caged BeKm-1 with an increase in inter-beat interval (648.7 ± 75.2 ms in illuminated condition *versus* 379.7 ± 13.8 ms in non-illuminated larvae, n= 10, p<0.05; Figure 5d) and episodes of atrio-ventricular block in 7/11 larvae (Figure 5e, Movie S3-4).

**Figure 5.**
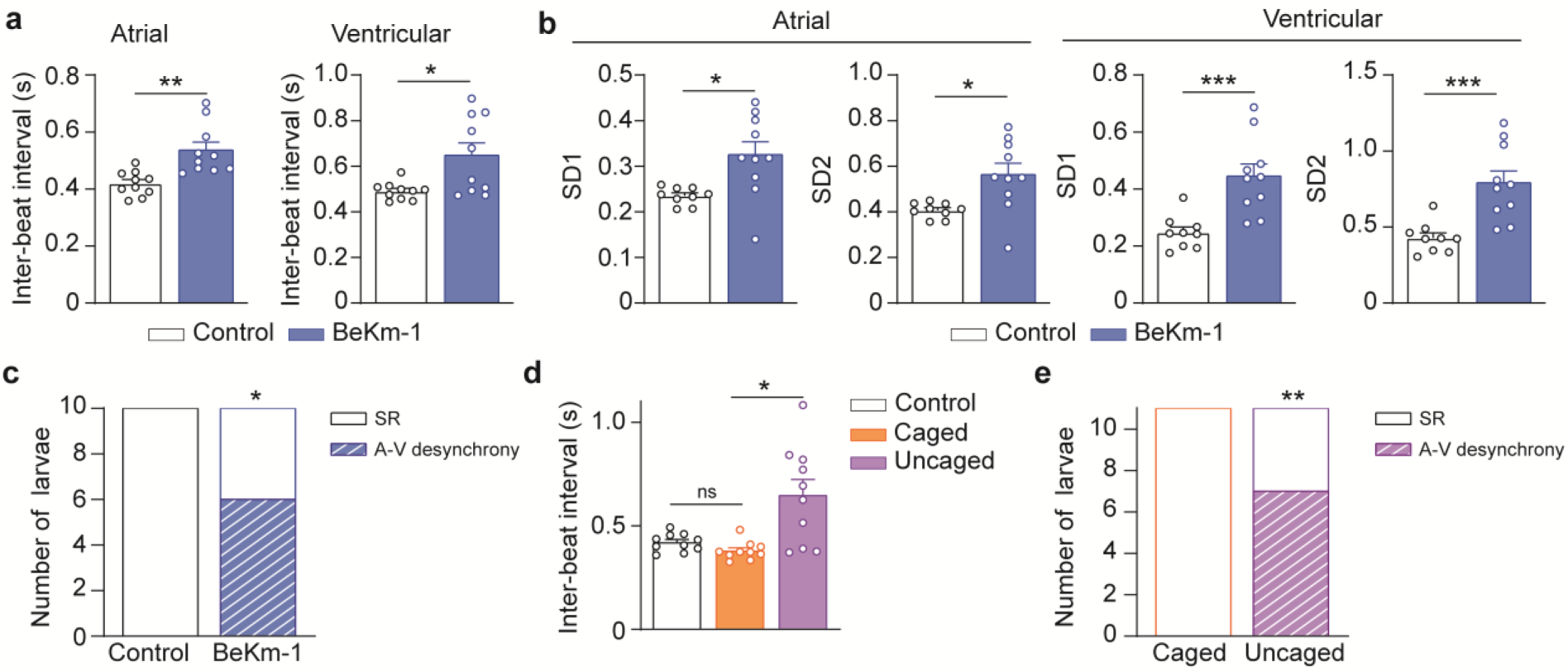
*In vivo* photopharmacological inhibition of ERG zebrafish larvae. (a) Injection of 50 µM BeKm-1 in zebrafish larvae produces both atrial (left) and ventricular bradycardia (atrial inter-beat interval of 530.7 ± 27.9 ms in the presence of BeKm-1 *versus* 420.2 ± 13.5 ms in control, n= 10 larvae, **, p<0.01; ventricular inter-beat interval of 648.8 ± 53.8 ms in the presence of BeKm-1 *versus* 491.2 ± 12.1 ms in control, n= 10 larvae, *p<0.05). The slight difference observed between atrial and ventricular inter-beat intervals is non-significant. (b) Injection of 50 µM BeKm-1 in zebrafish larvae increases both atrial (left, *, p<0.05) and ventricular (right, ***, p<0.001) rhythm heterogeneity as illustrated by increase in SD1 and SD2 coefficients. (c) Quantification of atrio-ventricular desynchronization in control conditions of after injection of 50 µM BeKm-1 (n = 10 larvae, Fisher test *p<0.05) based on video evidence (Movie S1-2). (d) Quantification of bradycardic effects of injection of 50 µM of caged BeKm-1 in zebrafish larvae (inter-beat interval of 379.7 ± 13.8 ms in caged BeKm-1 (orange) *versus* 420.2 ± 13.6 ms in control (black), n= 10 larvae, p=ns) and of uncaging in the same larvae (inter-beat interval: 648.7 ± 75.2 ms in illuminated condition *versus* 379.7 ± 13.8 ms in non-illuminated larvae, n = 10 larvae, *, p<0.05). (e) Quantification of atrio-ventricular desynchrony in control conditions of after injection of 50 µM caged BeKm-1 (n = 10 larvae, Fisher test *, p<0.05) based on video evidence (Movie S3-4). Data are presented as mean ± SEM.

### *In vivo* optical control of ERG activity in rats

We next validated our approach of photopharmacology control of heart rhythm in anesthetized rats undergoing right mini thoracotomy, which enabled visualization and illumination of the right atrium (Figure 6a). Concomitant administration of atropine and propranolol was applied similarly to experiments carried out in mice (Figure S1a). Intraperitoneal injection of 300 μg/kg caged BeKm-1 produced a minor effect on sinus rhythm with a ΔRR of 6.6 ± 1.5 ms (Figure 6b,c). A 10-min UV-illumination of the right atria with a 1-mm diameter light guide (365 nm, 75 mW/cm²) (Figure 6a) led to a final ΔRR of 15.9 ± 2.9 ms compared to control condition (Figure 6b,c). These results highlight for the first time that cardiac heart rhythm can be controlled experimentally in mammals by peptide-based photoactivation *in vivo*.

**Figure 6.**
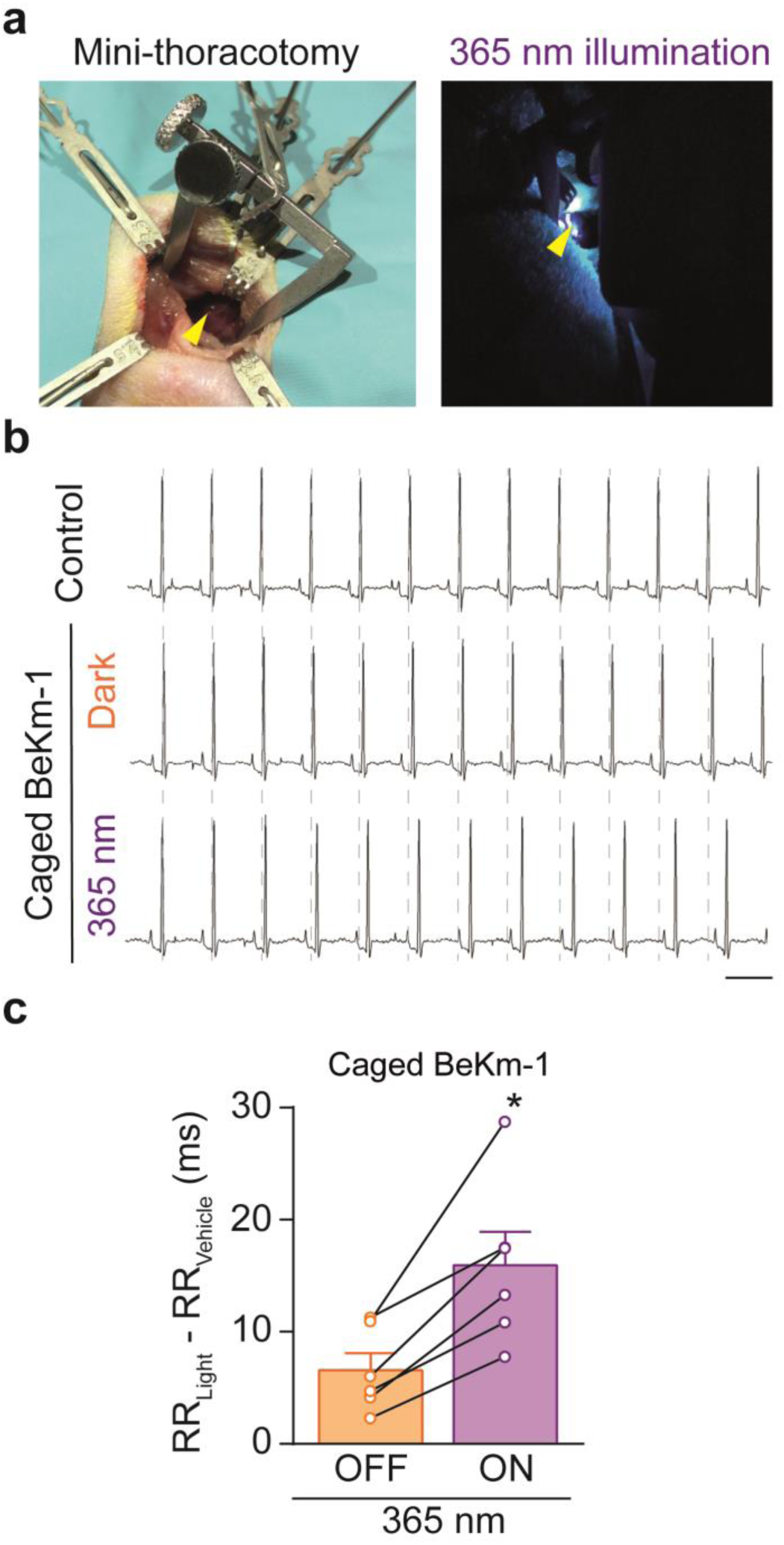
Photopharmacological bradycardia in anesthetized rat. (a) Photographs illustrating the right mini-thoracotomy procedure in rat (left) and the illumination area (right). The arrow indicates the right atrium. (b) Representative electrocardiograms recorded in control conditions (after thoracotomy), after injection of 5 µg/kg of caged BeKm-1 prior to 365 nm illumination (orange) and after 10 min of illumination (purple). Scale bar = 150 ms. (c) Quantification of increase in RR interval mediated by caged (orange) and uncaged (purple) BeKm-1 (caged ΔRR of 6.6 ± 1.5 ms *versus* uncaged ΔRR of 15.9 ± 2.9 ms, n = 6 rats, p<0.05).

## Discussion

In this study, we have demonstrated for the first time the applicability of photopharmacology for the control of heart electrical activity by using, in its caged form, a natural peptide active against the ERG channel. To our knowledge, there is no other example in the scientific literature detailing such an application. These data extend to the heart the possibility of using caged natural peptides for the control of excitability^18^. Using natural peptides derived from animal venoms offers a clear advantage for *in vivo* applications. They possess, for several of them, great selectivity and act at low concentrations. Thanks to evolutionary pressure, they possess long half-lives *in vivo* as well as enhanced targeting properties^45^. The fact that they are peptides provides several opportunities for chemical modifications, as described here for caging, but also for attaching cargoes^46^ or imaging modalities^47^. In the case of caged BeKm-1, 300 μg/kg was sufficient to achieve regulation of heart activity by light *in vivo* in rats. The chosen target, namely the ERG channel, was far from trivial for such a demonstration as this channel is poorly expressed in rodents. We therefore expect that even stronger demonstrations will be made with this technology by targeting more abundant ion channels in the future.

The choice of a natural peptide active against the ERG channel may seem surprising as ERG blockage is recognized for its pro-arrhythmic properties. However, the potential of the photoactivation technique is to limit the block of ERG channel activity to spatially-restricted cardiac areas. In this sense, it is possible to envision uncaging of the BeKm-1 peptide solely into the sino-atrial node to induce negative chronotropism and heart rate reduction in the absence of direct proarrhythmic effect - expected if the peptide was also active in the ventricle - in other preclinical models and in humans. While such an approach may seem counterintuitive at the scientific level considering the wealth of literature on the role of ERG channels in cardiac safety, the technology could easily be applied to other channels which can also regulate cardiac rhythm without being preoccupied by these medical concerns. Several other ion channels have been involved in the genesis and regulation of cardiac pacemaker activity: G protein activated K^+^ (GIRK)^48^, hyperpolarization activated f-(HCN)^49^, Ca_v_3.1^50^, Ca_v_1.3^35, 51^ Ca^2+^ and Na^+^ neuronal Na_v_1.1 channels^52^ to name a few. For these channel types, specific natural peptide blockers are not always available. Therefore, straightforward transposition of this technology to target these channels in the safest manner with full specificity will require further research. However, we are not fully devoid of solutions as several peptides have been identified for some of these channels: tertiapin Q for GIRK channels^53^ and huwentoxin IV for Na_v_1.1^18^. While tertiapin Q is a specific blocker for GIRK channels, huwentoxin IV lacks specificity towards Na_v_1.1 as it also targets other neuronal Na_v_ channels. Nevertheless, the advantage of the uncaging approach is that, by spatially-restricting the release of the active peptide in a given area of the tissue, an artificial form of channel type specificity can be reached by illumination. It is worth mentioning, in the context of the use of natural peptides, that channel activation is also possible through gating modulation^54^. This positive channel modulation opens interesting opposite perspectives such as accelerating heart rate in patients carrying sino-atrial node dysfunction Photoactivation of natural peptides can also be used in different contexts for basic science and preclinical studies. The most obvious application would be the definition (step 1) / delineation (step 2) of proarrhythmic cardiac areas with regard to the distribution of a given channel type. For instance, it would be of interest to dissect the contributing role of sodium channels in the arrhythmias initiated in the right ventricular outflow tract using this technology. A second less evident application, but clinically more relevant, would be the pharmacological correction of arrhythmias by light. Experimentally, optogenetics was proven successful in terminating ventricular arrhythmias induced by tachypacing^10, 11^. Conceptually, photopharmacology should also prove successful provided that the right channel types are targeted in the pro-arrhythmic region. For this purpose, several targets can be envisioned from K^+^ to Na^+^ channels offering a large choice of natural peptide compounds. Such a technology would be equivalent to radiofrequency ablation without permanent necrosis of the cardiac tissue targeted. It would however induce transient effect, which in some cases may be desirable at the clinical level.

It is important to note that photopharmacology aimed at regulating ion channel activity is only of real interest if the drug has a defined duration in its action on its target. For rhythm regulation, it would be interesting to prolong the mode of action of the released active peptide in a limited area if the target is expressed in whole heart (such as ERG channel), while for arrhythmia correction restricted duration of action would be beneficial. For these reasons, it is fortunate that natural peptides come into many flavors, some display a good action reversibility, while others are almost irreversible. Peptides being versatile chemical platforms, new chemistries may be developed/added with the intention to modulate their duration of action to perfectly match the clinical needs.

Of course, photoactivation of caged peptides has no future at the preclinical and clinical levels without the development of implantable devices. Fortunately, the field of wireless optical stimulators has been blooming and several important initiatives have emerged^55, 56^, including for the heart. Notably, these devices have proven successful in animal models without any inconvenience which illustrates also that the technology is mature enough to be translated to human hearts in a similar manner to pacemaker devices.

## Acknowledgements

We thank the animal facility care unit (UTE IRS-UN) for animal housing and management. The authors also thank the NMR sub-platform of the MO2VING facility (CBM, Orléans, France).

